# CRISPR/Cas9-mediated Generation of *COL7A1*-deficient Keratinocyte Model of Recessive Dystrophic Epidermolysis Bullosa

**DOI:** 10.1101/2023.06.15.545036

**Authors:** Farzad Alipour, Mana Ahmadraji, Elham Yektadoust, Parvaneh Mohammadi, Hossein Baharvand, Mohsen Basiri

## Abstract

**Objective:** Recessive dystrophic epidermolysis bullosa (RDEB) is a genetic skin fragility and ultimately lethal blistering disease caused by mutations in the *COL7A1* gene which is responsible for coding type VII collagen. Investigating the pathological mechanisms and novel candidate therapies for RDEB could be fostered by new cellular models. Here, we developed multiple immortalized *COL7A1*-deficient keratinocyte cell lines using CRISPR/Cas9 technology as RDEB cellular model.

**Materials and Methods:** In this experimental study, we used transient transfection to express *COL7A1*-targeting gRNA and Cas9 in HEK001 immortalized keratinocyte cell line followed by enrichment with fluorescent-activated cell sorting (FACS) via GFP expressing cells (GFP^+^ HEK001). Homogenous single-cell clones were then isolated, genotyped, and evaluated for type VII collagen expression. We performed a scratch assay to confirm the functional effect of *COL7A1* knockout.

**Results:** We achieved 46.1% (p < 0.001) efficiency of indel induction in the enriched transfected cell population. Except for 4% of single nucleotide insertions, the remaining indels were deletions of different sizes. Out of nine single clones expanded, two homozygous and two heterozygous *COL7A1*-deficient cell lines were obtained with defined mutation sequences. No off-target effect was detected in the knockout cell lines. Immunostaining and western blot analysis showed the lack of type VII collagen (COL7A1) protein expression in these cell lines. We also showed that *COL7A1*-deficient cells had higher motility compared with their wild-type counterparts.

**Conclusion:** We reported the first isogenic immortalized *COL7A1*-deficient keratinocyte lines that provide a useful cell culture model to investigate aspects of RDEB biology and potential therapeutic options.

## Introduction

Epidermolysis bullosa (EB) is a group of inherited skin blistering diseases, characterized by fragility of the skin due to defective epithelial cell adhesion leading to blister, erosion, and ulcers (1). In this group of diseases, 16 genes detected that their proteins in keratinocyte and fibroblast participate in the cell cytoskeleton and extracellular matrix. According to phenotypic heterogenicity, there are four main types including EB simplex (EBS), junctional EB (JEB), dystrophic EB (DEB), and Kindler syndrome (2). The severest form of these diseases is recessive dystrophic epidermolysis bullosa (RDEB), caused by loss of function mutations in *COL7A1*, the gene codes for type VII collagen protein (3). This protein is the main component of anchoring fibrils which is crucial for the adherence of the dermal-epidermal basement membrane. Mutations in *COL7A1* cause a variety of severities where truncated forms manifest a relatively milder forms and complete lack of the protein results in severest forms of RDEB (4).

Experimental studies on rare genetic disorders such as RDEB confront ethical and practical challenges including paucity of available stable patient donors, technical inconsistency in sampling and cell isolation, patient discomfort, and variability in the severity of disease between patients (5). Moreover, RDEB primary cells are mostly challenging due to inefficient and labor-consuming isolation, expansion, and limited proliferation capacity which shows the need for cell lines that easily be handled for experimental purposes (6). Although animal models are available for EB disorders, there are some shortenings such as imperfection in development, stillbirth, and deficiency in the recapitulation of disease phenotype (7). Therefore, there is a need to establish new cellular RDEB models to study pathological pathways and evaluate potential therapeutic approaches.

Gene editing technologies such as clustered regularly interspaced short palindromic repeats (CRISPR)/CRISPR associated protein 9 (Cas9) technology have provided a straightforward means to introduce mutations to normal cells for modeling purposes. Site-specific double-strand break (DSB) induced by CRISPR/Cas9 complex undergoes the error-prone non-homologous end joining (NHEJ) DNA repair, which may result in insertion-deletion (indel) mutations. A portion of indel mutations causes frameshift or other types of detrimental changes in the gene sequence (8). Introducing mutations to the disease-related genes in the context of a normal cell line has been used as a powerful method to generate isogenic cellular models for the disease (9-14).

In this study, we used CRISPR/Cas9 technology for the generation of immortalized *COL7A1-* deficient keratinocytes as a cellular model for RDEB. Since truncated *COL7A1* mutants represent mild manifestations of the disease and might hamper future investigations on exogenous type VII collagen therapeutics, we aimed to eliminate the expression of the hole protein by targeting the first exon. The resultant *COL7A1*-deficient keratinocyte cell lines were then characterized in terms of genotype and type VII collagen expression.

## Materials and Methods

### Guide RNA design and construction

To induce frameshift on the transcription sequence of *COL7A1* designed one guide RNA (ACTGCCTAGGATGACGCTG) targeting the first exon of *COL7A1* gene based on minimal off-target activity by CRISPOR online bioinformatic tool (15). Designed complementary sense (ghCOL7A1-S) and antisense (ghCOL7A1-A) oligonucleotides (Table S1) incubated to hybridize together and constitute a double-strand oligonucleotide. The hybridized double-stranded (ghCOL7A1-S + ghCOL7A1-A) DNA fragment cloned into the Esp3I (BsmBI) restriction sites of the LentiCRISPRv2GFP (Addgene plasmid # 82416) using a simultaneous digestion-ligation reaction as described previously (16). Briefly, a mixture of 10 pmol of the insert DNA fragment, 1 μg of LentiCRISPRv2GFP plasmid, 5 units of Esp3I enzyme, 1 unit of T4 DNA ligase, 1 μL of 10X FastDigest buffer, and 1 μL of 10X T4 DNA ligase buffer (all reagents from Thermo Fisher Scientific, Waltham, MA, USA) in a total volume of 20 μL was incubated in 37°C for 3 hours. The reaction product containing the recombinant plasmid (LentiCRISPRv2GFP-ghCOL7A1) was transformed into chemo-competent *E. coli* cells. Restriction mapping and sequencing were done on single clones for verification of the correct gRNA sequence in the vector.

### Cell culture

Immortalized keratinocyte (HEK001) cell line (CRL-2404, ATCC) cultured on DMEM/F12 medium (Cat. No. 21331020, Gibco, Thermo Fisher Scientific), containing 10% (v/v) fetal bovine serum (Cat. No. 10270-106, Gibco, Thermo Fisher Scientific), 2 mM GlutaMAX (Cat. No. 35050038, Gibco, Thermo Fisher Scientific), 1% (v/v) NEAA (Cat. No. 11140050, Gibco, Thermo Fisher Scientific), 1% (v/v) PenStrep (Cat. No. 15140122, Gibco, Thermo Fisher Scientific) and supplemented with KGM-GoldsingleQuots (Cat. No. 001922152, Lonza, Switzerland), containing Bovine pituitary extract (BPE), insulin (5 mg/mL), epidermal growth factor (EGF, 10 ng/mL), epinephrine, hydrocortisone (0.4 mg/ml), transferrin (17). The cells were incubated at 37°C in a 5% (vol/vol) CO_2_ incubator over the experiments.

### Transfection of HEK001 cells

HEK001 cells were cultured in a 100-mm culture dish to reach 70-90% confluency. Two to three hours before transfection, the media was replaced with a fresh complete medium without antibiotics. HEK001 cells were transfected with LentiCRISPRv2GFP-ghCOL7A1 plasmid using lipofectamine 2000 Transfection Reagent (Cat. No. 11668019, Thermo Fisher Scientific) according to the manufacturer’s instructions. Briefly, a transfection medium was prepared by adding 24 μg of vector and 24 μl of lipofectamine 2000 to 1 ml base medium of DMEM/F12 in two separate tubes followed by 10 min incubation. The vector-containing medium was added to the lipofectamine tube and incubated for another 15 min. Transfection medium was added to cells gently and incubated overnight. The next day, the medium was replaced with a fresh keratinocyte growth medium and grown for another 36 hours. All Incubation was done at 37°C in a humidified 5% (vol/vol) CO_2_ incubator.

### Single-cell clone isolation

Since in our hand, single HEK001 cells had reduced viability right after FACS isolation we used a limited dilution method (18, 19) to obtain single-cell clones. Two days after transfection by LentiCRISPRv2GFP-ghCOL7A1 plasmid, around 2×10^5^ HEK001 cells expressed GFP (GFP^+^ HEK001) isolated by FACS Aria II cell sorter (BD Biosciences, San Jose, CA, USA) and cultured in one well of a 24-well plate. When 70-90% confluency was reached, the cells were detached by trypsin (Cat. No. 25200056, Gibco, Thermo Fisher Scientific), counted, and serially diluted in complete media to obtain a cell suspension of 5 cells/ml. Then, 100 μl of this suspension was manually seeded in each well of a 96-well plate to obtain an average of 0.5 cells per well. The wells containing truly single cells were determined using an inverted microscope (Olympus IX71S1F-3, Japan). Selected single-cell clones then were serially expanded into 48-, 24-, 12-, and 6-well plates for downstream analyses.

### Sanger sequencing and targeting efficiency assessment

Genome extraction was done with a commercial genome extraction kit (Cat. No. FABGK 100, Favorgen Biotech Corporation, Taiwan) according to the manufacturer’s instructions. Briefly, 10^6^ cells were resuspended in the lysis buffer of the kit and incubated for 10 min to obtain clear cell lysate. Genomic DNA was isolated from the cell lysate using silica columns and washing buffers provided in the kit. The gRNA target region was amplified by polymerase chain reaction (PCR) using COL7A1-F1 and COL7A1-R1 primers (Table S1) with the following PCR condition: 95°C for 10 min, 35 cycles of 95°C for 30 s, 60°C for 30 s, 72°C for 45 s and then 72°C for 10 min. The PCR product was subjected to Sanger sequencing using COL7A1-F2 and COL7A1-R2 primers (Table S1). Sequencing data were analyzed for insertion/deletion events and targeting efficiency using the Tracking of Indels by Decomposition (TIDE) analysis method (20).

### Off-target prediction and sequencing

Potential off-target sites in the human genome were predicted using CRISPOR bioinformatic tool (15). Four off-target sites in the exonic regions of *FAT3, ANKZF1, AXIN2*, and *NPHP4* genes were evaluated with PCR and sequencing. PCR program was done as follows: 95°C for 10 min, 35 cycles of 95°C for 30 s, 60°C for 30 s, and 72°C for 45 s, followed by a final extension step at 72°C for 10 min). PCR products were subjected to Sanger sequencing. Primers used for the PCR and sequencing reactions are listed in Table S1.

### Immunofluorescence staining

The cells were fixed with paraformaldehyde 4% at 4°C for 20 minutes, washed three times with phosphate buffer saline (PBS) containing 0.05% Tween 20 (PBST, Cat. No. P2287-100ML, Sigma), and then permeabilized with 0.5% Triton X-100 (Cat. No. X100-1L, Sigma) at room temperature for 10 minutes. Blocking was performed with goat serum 10% at 37°C for 60 min and incubated with rabbit polyclonal to type VII collagen (ab93350, Abcam, Cambridge, UK) at 1:100 overnight at 4°C. The following day, cells were incubated with Alexa Fluor 546-conjugated goat anti-rabbit secondary antibody, (A-11035, Thermo Fisher Scientific) at 1:500 for 45 minutes at room temperature, and subsequently, nuclei were stained with DAPI 20 mg/ml (D8417, Sigma). The cells were analyzed by Olympus IX71S1F-3 inverted fluorescence microscope (Olympus, Tokyo, Japan) equipped with a 100 W Mercury light source (Olympus U-LH100HG) as a source of excitation radiation and Olympus U-TV0.63XC image acquisition system. The staining observed by 10× objective lenses for GFP (excitation 460-490 nm, emission 520 nm, exposure time 500 ms), Alexa Four546 (excitation 510-555 nm, Emission 590 nm, exposure time 500ms) in exposure time (500 ms) and DAPI (excitation 330-385 nm, emission 420 nm, exposure time 10 ms).

### Western blot analysis

To extract total protein, cell pellets were lysed by adding cell lysis buffer (7 M urea, 2 M thiourea, 4% CHAPS, 30 mM Tris, 40 mM dithiothreitol) and 1X cocktail protease inhibitor, vortexing for 5 min on ice, sonicating for 5 min on ice, followed by centrifuging at 14000 g for 15 min and transferring the supernatant to new tubes. Protein concentration measured by pierce BCA assay (Cat No. 23227, Thermo Fisher Scientific). Protein samples were denatured at 50°C for 60 min in 50 μl 5X loading buffer (Laemmle 5X buffer: 0.5 M Tris-HCl, pH 6.8, 10% glycerol, 2% SDS, 0.025% bromophenol blue and 50 mM dithiothreitol), then separated by loading on 8% sodium dodecyl sulfate-polyacrylamide gel electrophoresis (SDS-PAGE), 100 V, 2 hrs., followed by electrotransfer to polyvinylidene fluoride membranes at 15 V at 15°C, overnight. Membranes were blocked with 2% BSA in Tris-buffered saline containing 0.05% Tween (TBST) 20 at room temperature for 2 hrs, followed by adding primary antibody rabbit anti-COL7A1 (ab93350, Abcam) at 1:1000 in TBST and incubated overnight. A polyclonal β-tubulin antibody (sc15335, Santa Cruz) at 1:1000 served as loading control. The next day, after washing three times in TBST 0.1%, the blots were incubated at room temperature for 1 hr. with HRP-conjugated donkey anti-rabbit secondary antibody at 1:50000 in TBST, followed by three washes. To detect the desired band, SuperSignal West Femto Maximum Sensitivity substrate (Cat No. 34096, Thermo Scientific) was added, and Chemiluminescent signals were captured by the Alliance Q9 Advanced Uvitec System.

### Statistical analysis

TIDE analysis uses a built-in two-tailed t-test of the variance-covariance matrix of the standard errors to calculate p-values for the frequency of each indel mutation (20). To compare the normalized percentage of cell-free area in the scratch assay, we performed a two-way ANOVA followed by Sidak’s multiple comparison test using GraphPad Prism 8.0.2 (GraphPad Software Inc., La Jolla, CA, USA). The adjusted p-values less than 0.01 were considered statistically significant.

### Ethics approval

The study was approved by the E Ethical Committee of Royan Institute - Academic Center for Education, Culture, and Research (IR.ACECR.ROYAN.REC.1398.076).

## Results

### Guide RNA Design and generation of *COL7A1*-knockout HEK001 cells

To knock out the *COL7A1* gene in HEK001 immortalized keratinocytes, we designed a gRNA targeting the first exon of the gene. Corresponding sequence subcloned under a U6 promoter in an expression plasmid vector containing Cas9 coding sequence linked to GFP through a self-splitting 2A linker (Fig. 1A and B). Transfection of HEK001 cells by this vector resulted in a population of GFP^+^ cells with a transduction efficiency of 14.1 % (Fig. 1C). To enrich the gene-modified cells GFP^+^ cells were isolated by FACS (Fig. 1D).

**Fig. 1:**
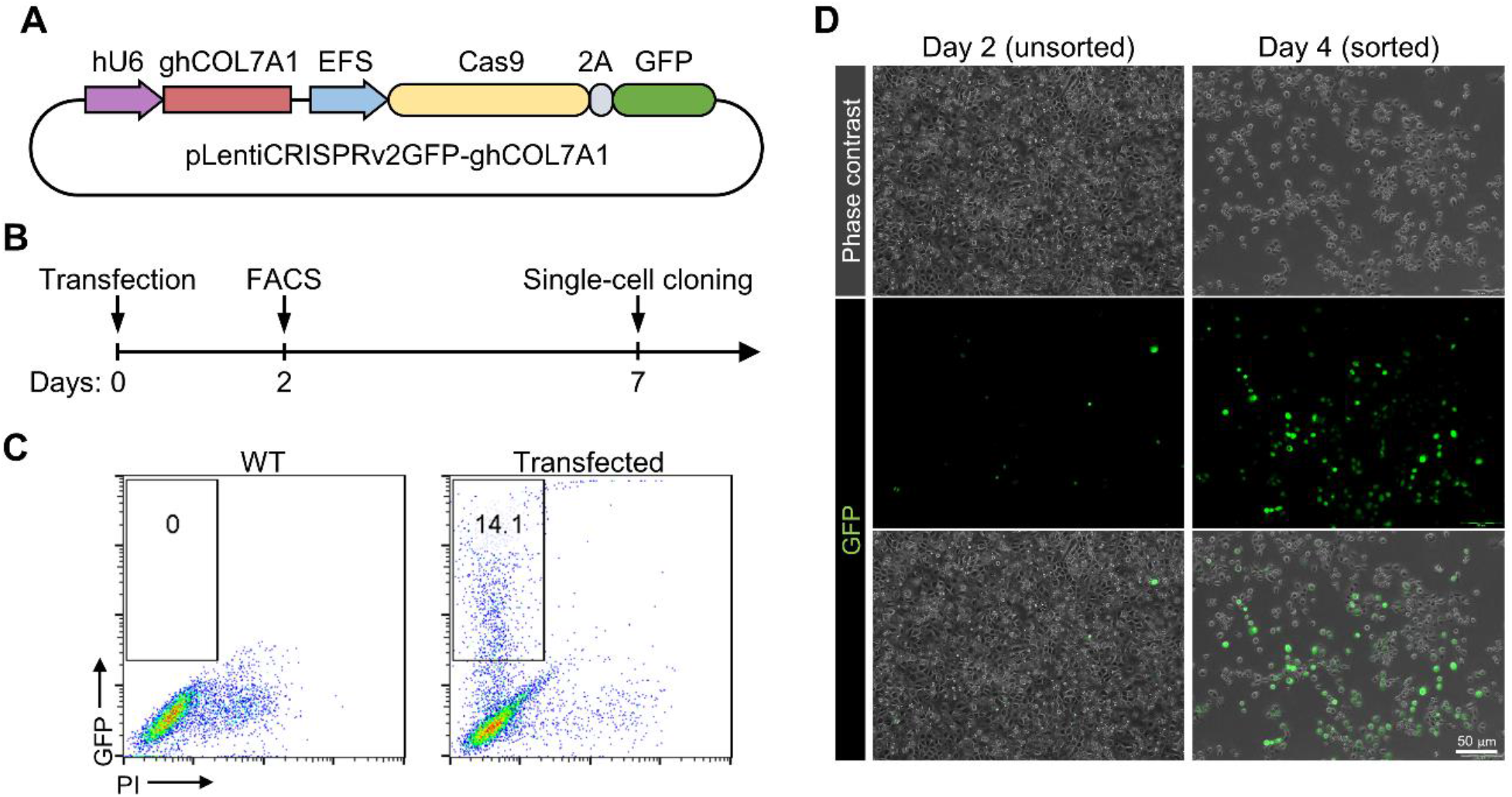
Transfection and isolation of GFP^+^ cells. (A) Schematic map of the CRISPR plasmid vector. gRNA insert in expression vector encompassing Cas9 and GFP sequence under U6 promoter. (B) Diagram of the experimental procedure showing transfection, enrichment by fluorescence-activated cell sorting (FACS), and single-cell cloning by limited dilution. (C) Viable GFP-expressing cells comprising 14% of the transfected keratinocytes were sorted to enrich the genome-edited cell population. (D) Fluorescent microscopy of the pool population of cells 2 days after transfection (before sorting), and enriched GFP^+^ cells 2 days after sorting.

### Sequencing analysis of the pool population of transfected cells

To assess targeting efficiency in transfected HEK001 cells, genomic DNA of the isolated GFP^+^ cells was extracted and PCR amplicon spanning gRNA targeted site was sequenced by Sanger sequencing. Results showed a composite sequence trace after the break site resulted from the presence of indel mutations in the pool population of transfected cells (Fig. 2A). Decomposition of the sequence trace data (Fig. 2B) showed that the overall editing (indel) efficiency was 46.1% (*p* < 0.001) with more frequent deletion events compare to nucleotide insertions (Fig. 2C). We did not detect insertions of more than one nucleotide in the bulk population. Single nucleotide insertion was observed in 4.0% (*p* < 0.001) of sequences containing either T or A nucleotide insertions (Fig. 2D).

**Fig. 2:**
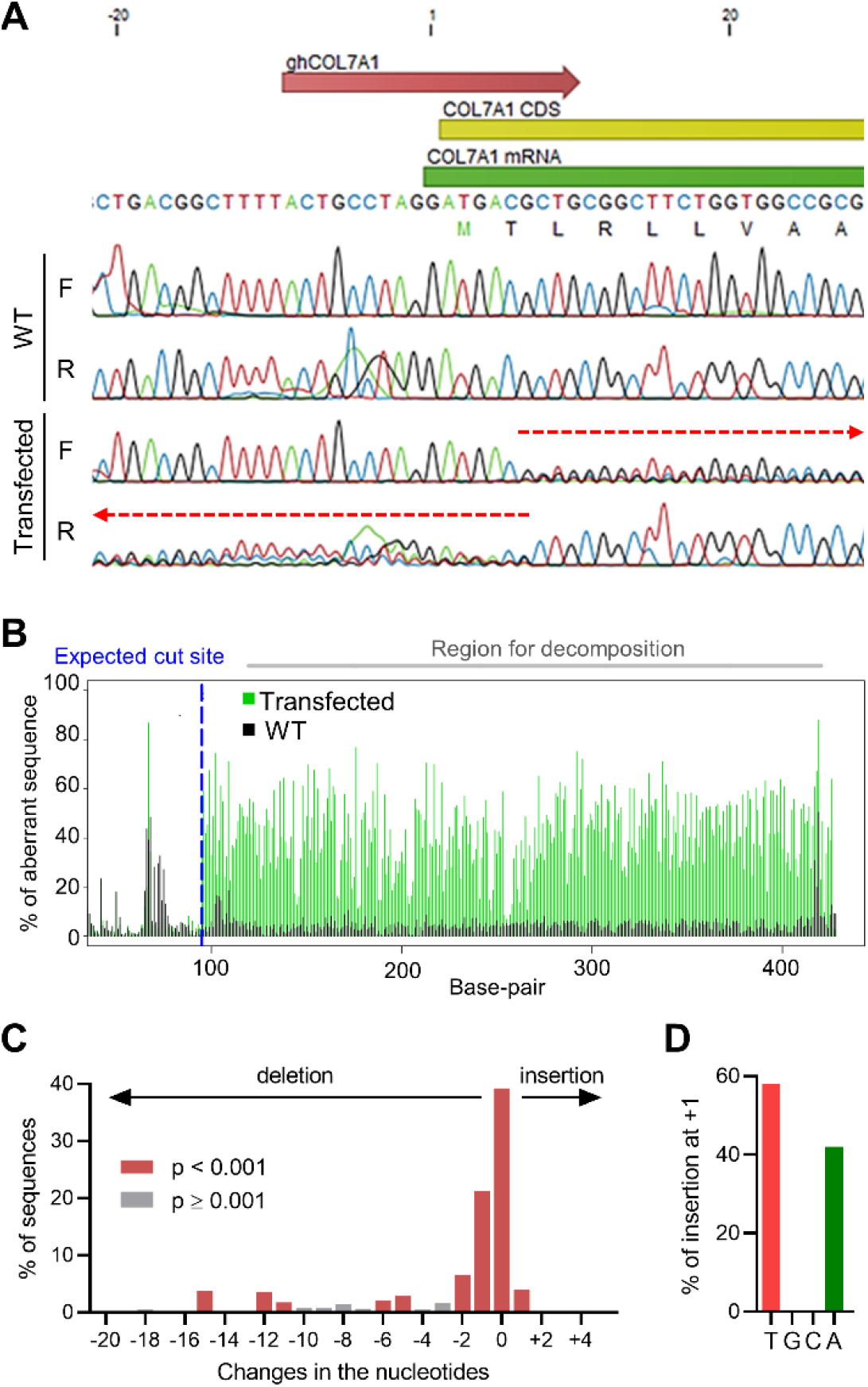
Sanger sequencing and decomposition analysis on the bulk population of transfected HEK001 cells. (A) Sequencing on the bulk population of transfected cells represents a composite sequence trace after the break site based on the direction of the primer. (B) Comparison of transfected bulk with untransfected revealed aberrant sequence after the break site. (C) Decomposition analysis of changes in the targeted region shows the frequency of mutations based on the number of nucleotides deleted from or inserted into the sequence. (D) Out of all single nucleotide insertions, 58% was T and 42% was A. F: forward primer, R: reverse primer, WT: wild-type.

### Sequencing analysis of the single-cell clones

To establish a homogeneous cell line with identified sequence, we isolated single-cell clones from the bulk transfected sequence. Out of 25 clones initially obtained from two 96-well plates, only nine showed normal expansion and were subjected to further molecular analysis. Decomposition analysis of the target genomic sequence in these nine clones (Fig. 3A) showed that one clone (B6) only contains the intact wild-type allele while in three clones (B2, O3, and T7) more than two alleles were indicated. Clone B1 showed a heterozygous genotype with single nucleotide insertion in one allele and an in-frame 15-nucleotide deletion in the other. The above-mentioned clones were excluded from the study due to the presence of wild-type sequences, in-frame mutations, or possible heterogeneity. However, four other clones (namely clones B4, T5, T6, and T8) showed genotypes compatible with the complete abolition of the *COL7A1* function. Clones B4 and T8 had heterozygous genotypes with deleterious frameshift mutations in both alleles (Fig. 3B and C). A single-nucleotide deletion at positions +9 (ninth nucleotide after the transcription start site) and a two-nucleotide deletion at positions +7 to +8 in clone B4 and a single-nucleotide deletion at position +7 in clone T8 introduce frameshifts into the *COL7A1* coding sequence which results in altered amino acid sequence and ectopic stop codons. The second allele of clone T8 had a 7-nucleotide deletion (positions +1 to +7) eliminating the start codon. Clones T5 and T6 showed homozygous deletion of, respectively, 33 nucleotides (positions −26 to +8) and 32 nucleotides (positions −21 to +12) encompassing both the transcription start site and start codon of *COL7A1* gene (Fig. 3C).

**Fig. 3:**
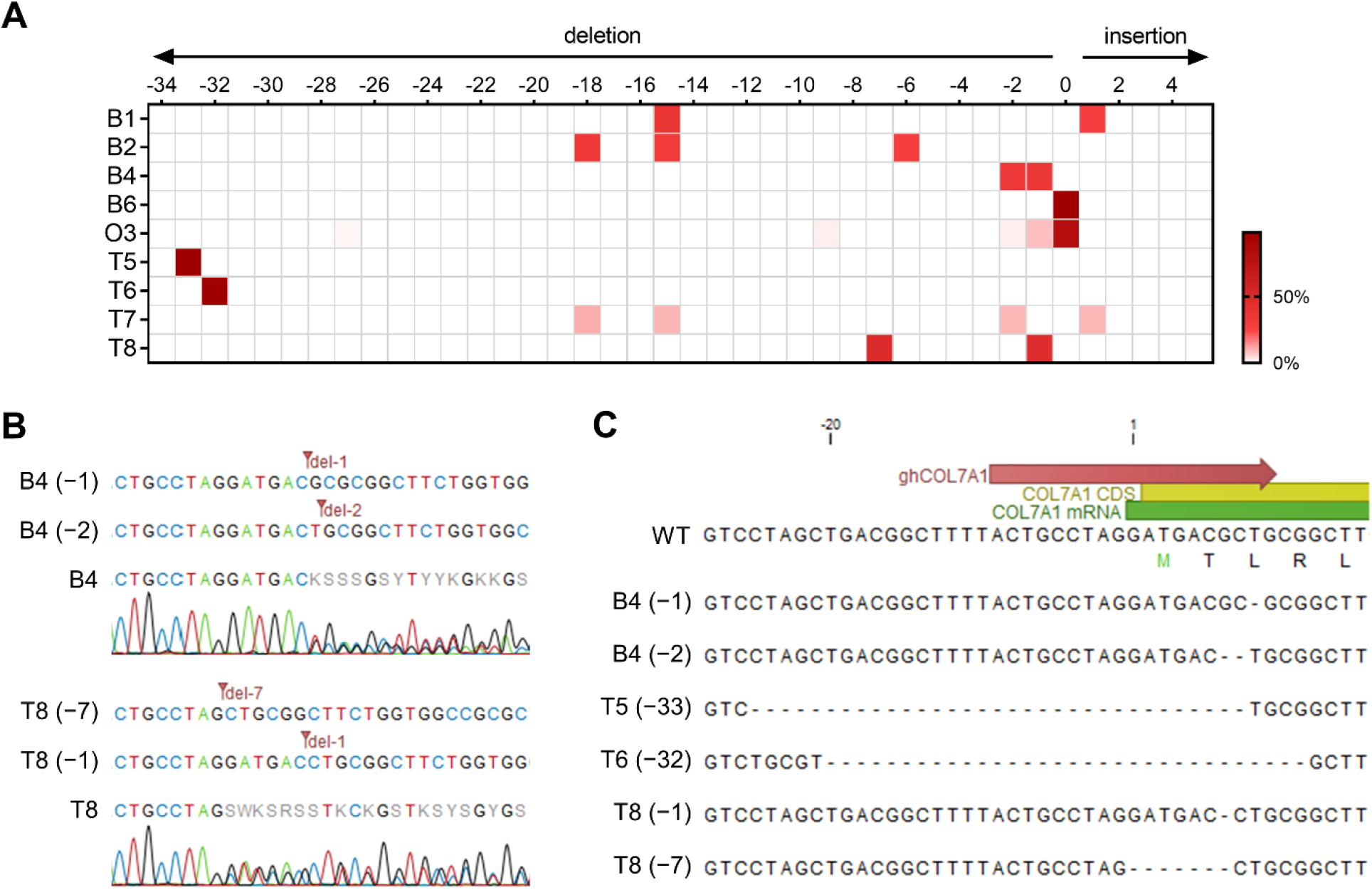
Genotyping of the single-cell clones. (A) Heatmap summarizes the results of decomposition analysis on the sequences data obtained from nine single-cell clones shown on the vertical axis. Only the genotypes with statistically significant (*p* < 0.001) frequency compared with the control sequence are depicted. (B) Decomposition of alleles in two heterozygous clones (B4 and T8) with deleterious mutations (C) Deleterious alleles detected in four knockout clones with frameshift mutations in B4(−1), B4(−2)

### Off-target analysis

To ensure that selected clones do not contain undesirable mutations due to the off-target activity of the CRISPR system, we performed an in-silico off-target prediction. Among 70 predicted off-target sites with four or fewer mismatched nucleotides (Table S2), only four were located in exonic sequences of *FAT3, ANKZF1, AXIN2*, and *NPHP4* genes, hence carrying the risk of unwanted mutations in a coding sequence. However, PCR-sequencing of these potential exonic off-target sites in the *COL7A1*-knockout cell lines showed no mutation when compared with the wild-type sequence (Fig. 4).

**Fig. 4:**
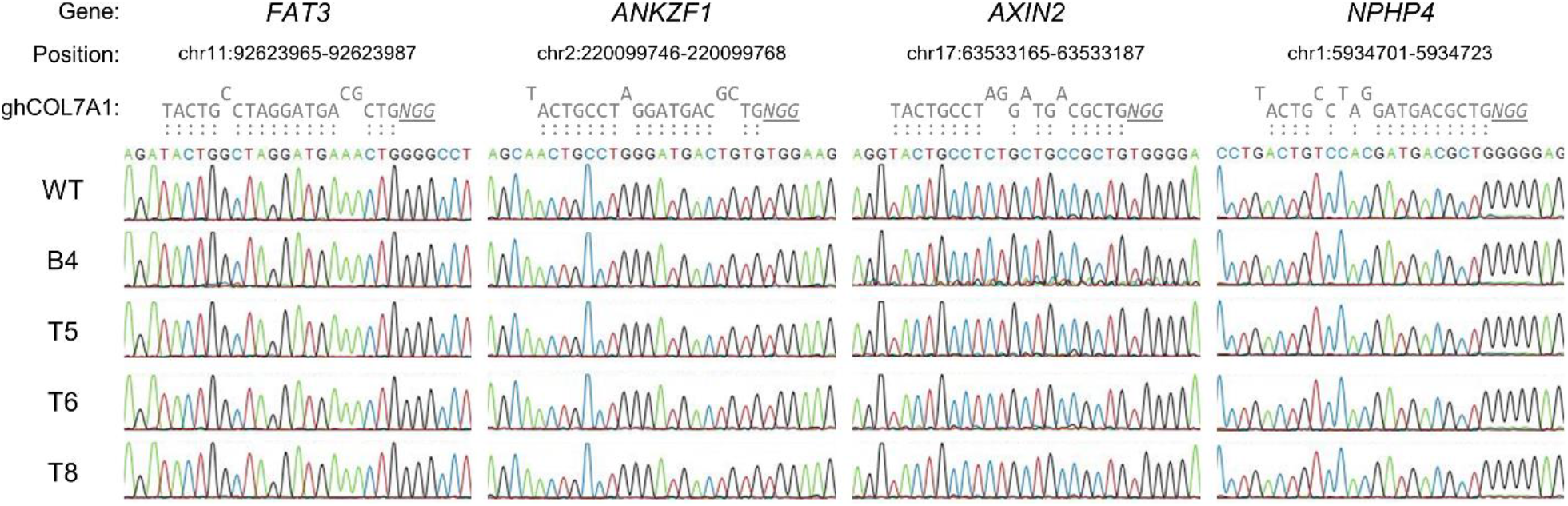
Investigating exonic potential off-target sites of ghCOL7A1 in knockout cell lines. Four predicted off-target sites in exonic sequences of FAT3, ANKZF1, AXIN2, and NPHP4 genes were amplified by PCR and subjected to DNA sequencing. Matched and mismatched nucleotides in comparison with ghCOL7A1 are shown above each sequence. No mutation was detected in any of these potential off-target sites.

**Fig. 5:**
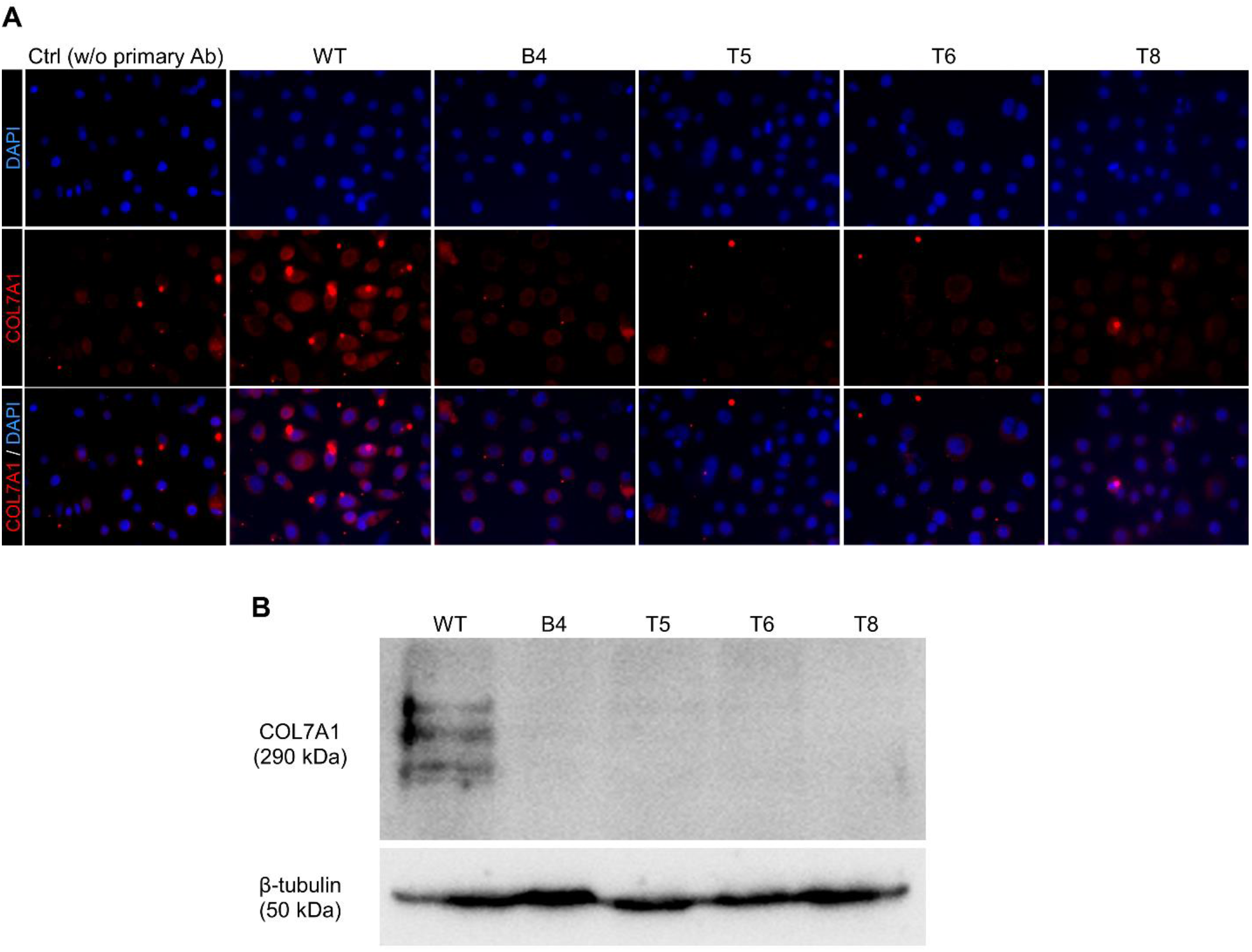
Evaluation of COL7A1 protein expression in the *COL7A1*-knockout HEK001 cell lines. (A) Four knockout cell lines (B4, T5, T6, and T8) were stained with type VII collagen (COL7A1)-specific antibody along with wild-type (WT) HEK001 cells. COL7A1 protein expression was not detected in any of the knockout cell lines as compared with the WT cells. A representative negative control without primary antibody (Ctrl w/o primary Ab) was included for comparison. (B) Western blot analysis of COL7A1 protein expression in the wild-type and knockout HEK001 cell lines. Human β-tubulin was stained as an internal control.

### Type VII collagen protein expression in selected knockout clones

Immunofluorescent staining was used to confirm the *COL7A1*-KO cell lines at the protein level. No expression of type VII collagen (COL7A1) was observed in the four clones selected in genotyping in comparison with wild-type HEK001 keratinocytes in immunostaining and Western blot analysis results (Fig. 4A and B). These results verified *COL7A1* deficiency predicted at sequence level on the four selected cell lines.

### Evaluation of cellular motility in the knockout cell lines

Since the loss of COL7A1 deficiency is associated with increased cellular motility, and elevated risk of keratinocyte malignancy and invasiveness in RDEB patients (21-23), we sought to measure cell motility in our *COL7A1*-KO HEK001 model as a functional indicate of *COL7A1* gene disruption. Using a scratch assay, we observed that *COL7A1*-KO HEK001 cells exert significantly higher migration compared with wild-type cells (Fig. 6 A and B).

**Fig. 6:**
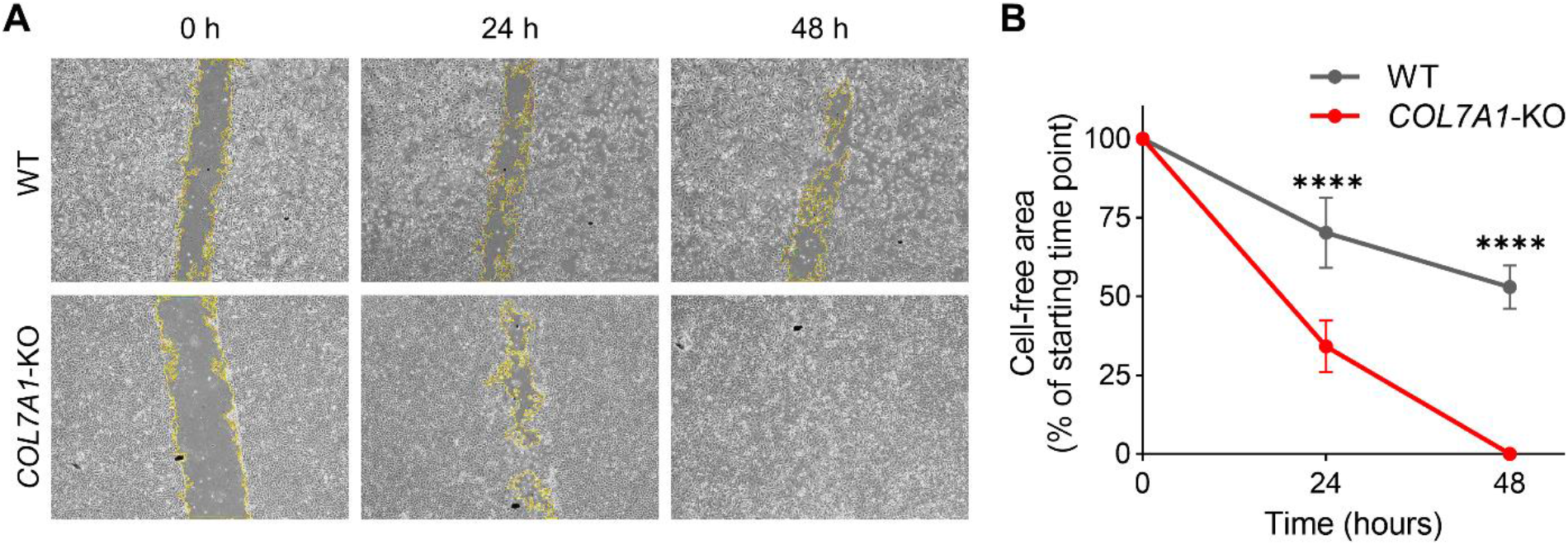
Assessment of cell motility by scratch assay in wild-type (WT) and *COL7A1*-KO cells. (A) Representative microscopic images of WT and one of the *COL7A1*-KO HEK001 cell lines (T6) at 0, 24, and 48 hours after wounding. (B) Quantitative comparison of cell migration and wound closure in WT and *COL7A1*-KO groups showed that *COL7A1*-KO HEK001 cells had obtained higher cell motility. Data presented as a percentage of the cell-free area of the starting time point ± SD; n = 3; **** p < 0.0001.

## Discussion

Investigation of rare diseases like EB has challenges with the availability of the proper patients and ethical issues as well as practical difficulties such as sampling, primary cell culture, and division potency of disease-specific cells. To obviate these problems, genetic modification in the desired gene to generate the disease model is favorable. In this study, we generated four immortalized keratinocyte cell lines harboring different characterized mutations of the *COL7A1* gene which can be used as a cellular model of RDEB.

The human *COL7A1* gene spans through more than 31 kb of chromosome 3 with 118 exons encoding a long polypeptide (around 2,944 aa) that gives rise to type VII collagen (24). Hundreds of *COL7A1* mutations have been detected in RDEB patients resulting in a variety of clinical manifestations with most severe forms correlating with the complete absence of the whole protein (25). Although amino acid substitution and truncated mutants can be found in some patients, these forms of the protein might be detected by antibody-based assays and therefore interferes with experimental investigations. Therefore, to generate a straightforward experimental tool, we chose to completely eliminate type VII collagen expression by targeting the first exon in the proximity of the start codon and the transcription start site. The results showed that this strategy successfully resulted in the generation of multiple mutations in four generated cell lines, however, we showed that all four selected cell lines lacked the expression of COL7A1 protein. In the functional level increased motility of the *COL7A1*-knocout cells is in line with previous findings showing that RDEB keratinocytes have higher motility associated with increased susceptibility to keratinocyte malignancies in RDEB patients (21-23).

*COL7A1* gene is expressed in both dermal fibroblasts and keratinocytes, with the latter being responsible for the majority of type VII collagen expressed in the skin (26). Therefore, several investigational cellular and molecular therapeutic strategies targeted keratinocytes in RDEB. Patient-derived immortalized keratinocytes have been generated as a cellular model for EB simplex (27) and RDEB (6). Although valuable for several studies, these cell lines lack the isogenic healthy counterparts. One disadvantage of patient-derived cell models compared to our approach is that they lack an isogenic control. Immortalized keratinocyte cell lines, derived from healthy donors, especially the HEK001 cell line, have been extensively characterized and used as valuable models for studying keratinocytes (28, 29). On the other hand, gene editing technologies such as CRISPR/Cas9 can be readily applied to generate targeted mutations in human cells. This approach has been also used in skin-related studies to investigate several biological functions such as keratinocyte development and homeostasis (30), signal transduction (31), adhesion (32), and epithelial differentiation (33, 34). In line with this approach, we employed CRISPR/Cas9 technology to generate new *COL7A1*-deficient HEK001 immortalized keratinocytes which closely resemble RDEB keratinocytes.

Although mouse models of RDEB have been generated and used for in vivo studies, rodent keratinocytes do not fully resemble human counterparts in terms of molecular mechanisms involved in EB, such as apoptotic and inflammatory pathways (35, 36). Targeting *COL7A1* by CRISPR/Cas9 in human primary keratinocytes has also been reported as a method to generate RDEB cellular model (7, 37). Whilst this method has the advantage of using primary keratinocytes, it also has potential limitations including the limited lifespan of primary keratinocytes, relatively labor-intensive procedure, generation of heterogenous mutations, persistent expression of gRNA and Cas9 by the integrative lentiviral vector, and the possibility of off-target mutations. Additionally, the efficiency of *COL7A1* knockout using CRISPR/Cas9 nucleoprotein in primary human keratinocytes was reported around 40% (8), while our approach allowed for the generation of homogenous *COL7A1* knockout cell lines. On the other hand, the major limitation of our approach is that immortalized keratinocytes do not fully represent some keratinocyte characteristics (e.g. differentiation and proliferation) which should be considered for future application of this cellular model. Moreover, immortalized keratinocytes might not be suitable for some in vivo experiments due to their constant proliferation. However, despite its inherent limitations, our strategy provided a more accessible, robust, and homogenous cellular model that may address some of the shortcomings in other cellular and animal models.

## Conclusion

In this study, CRISPR/Cas9 technology was used to generate a keratinocyte model cell line for RDEB. The generated model cell lines not only can be used in experimental studies such as investigational cell and gene therapies but also can potentially be applied to high-throughput screening of candidate drugs for symptomatic treatments such as wound healing, reduction in blister numbers, as well as amelioration of itch and pain. Moreover, our experimental strategy for CRISPR/Cas9-mediated gene editing in keratinocyte cell lines can be extended to the genes involved in other types of EB as well as the characterization and targeted modulation of pathogenic molecular cascades.

## Acknowledgments

This work was supported by the funds from Royan Institute. The authors indicated no potential conflicts of interest.

## Authors’ Contributions

F.A., M.B.; Contributed to the design, implementation of the research, data analysis, and writing the manuscript. F.A., M.A., E.Y.; Helped with carrying out the experiments. H.B., P.M.; Provided critical feedback and helped to finalize the manuscript. All the authors read and approved the final manuscript.

