## Supplementary Tables for "CRISPR/Cas9-mediated Generation of *COL7A1*-deficient Keratinocyte Model of Recessive Dystrophic Epidermolysis Bullosa"

**Supplementary Table S1.** Oligonucleotides and primers

| Application | Oligonucleotide/primer | Sequence (5' → 3') |
|---|---|---|
| Generation of gRNA-coding fragment | ghCOL7A1-S | CACCGACTGCCTAGGATGACGCTG |
| | ghCOL7A1-A | AAACCAGCGTCATCCTAGGCAGTC |
| PCR amplification of gRNA target region | COL7A1-F1 | TGGATTGGCATAAGGCAGGG |
| | COL7A1-R2 | ATCTGCCCATGCAGAACCTT |
| Sequencing of the gRNA target region | COL7A1-F2 | CACCTCTCTCCC-TGTGCT |
| | COL7A1-R2 | ACGCGCAGGCAAGACCAG |
| PCR amplification and sequencing of *FAT3* off-target region | FAT3-F | CTTCCCTGCTCCTTTCCCAG |
| | FAT3-R | CAAGGACAGAGACACGCTGT |
| PCR amplification and sequencing of *ANKZF1* off-target region | ANKZF1-F | GTTGACTGTGGGGACTCTGG |
| | ANKZF1-R | CCTCCTGCCTTTCCCAACAT |
| PCR amplification and sequencing of *AXIN2* off-target region | AXIN2-F | AGGGGTCAGTGCCAAAACAT |
| | AXIN2-R | ACTACATCCACCACCATGCC |
| PCR amplification and sequencing of *NPHP4* off-target region | NPHP4-F | CACCCTTGAAGTCCCTCCAC |
| | NPHP4-R | GTCCTCTTCTGAGATCGCGG |

**Supplementary Table S2.** Predicted off-target sites. Exonic sites are highlighted.

| Off-target sequence | Mismatch position | Mismatch count | MIT off-target score | CDF off-target score | Chromosome position (hg19) | Locus | Genomic context |
|---|---|---|---|---|---|---|---|
| TAATGCCTAGGATGAGGCTGAAG | ..*............*.... | 2 | 0.37468 | 0.034567901 | chr9:117192882-117192904 | *DFNB31* | intron |
| TAATGCCTAGGATGACACTTGGG | ..*.............*..* | 3 | 0.530449741 | 0.566222222 | chr3:113151837-113151859 | *WDR52-AS1-WDR52/WDR52-AS1* | intergenic |
| TACTGGCTAGGATGAAACTGGGG | .....*.........**... | 3 | 0.112770207 | 0.466666667 | chr11:92623965-92623987 | *FAT3* | **exon** |
| TCCTGCCAAGGATGACGCTCCGG | .*.....*...........* | 3 | 1.492090395 | 0.269387755 | chr1:1068727-1068749 | *C1orf159-RP11-465B22.5* | intergenic |
| CACTGCCTGGGCTGACGCTGAGG | *.......*..*........ | 3 | 0.846167111 | 0.15037594 | chr20:61012678-61012700 | *RP5-908M14.5-GATA5* | intergenic |
| TCCTGCCAAGGTTGACGCTGAGA | .*.....*...*........ | 3 | 0.276977778 | 0.014550264 | chr18:66654718-66654740 | *RP11-861L17.3-RP11-861L17.2* | intergenic |
| CACTCCCCAGGATGAAGCTGTGG | *...*..*.......*.... | 4 | 0.272333333 | 0.673469388 | chr7:43183211-43183233 | *HECW1-IT1* | intron |
| TTCTACCCAGGATGAAGCTGTGG | .*..*..*.......*.... | 4 | 0.258544304 | 0.63030303 | chr7:157381590-157381612 | *PTPRN2* | intron |
| TTCTGCCAAGGATGAAACTGAGG | .*.....*.......**... | 4 | 0.104848333 | 0.543030303 | chr22:31646772-31646794 | *LIMK2* | intron |
| TCATACCTAGGATGGCGCTGGGG | .**.*.........*..... | 4 | 0.397208228 | 0.383603175 | chr4:28937913-28937935 | *MESTP3-RP11-292B1.2* | intergenic |
| GACTGTCCACGATGACGCTGGGG | *....*.*.*.......... | 4 | 0.797205949 | 0.355279503 | chr1:5934701-5934723 | *NPHP4* | **exon** |
| TACTGCTTTGGAAGAAGCTGAGG | ......*.*...*..*.... | 4 | 0.039742653 | 0.3375 | chr16:64250429-64250451 | *RP11-744D14.2-AC012322.1* | intergenic |
| TGCTGCCTTGGAAGACACTGAGG | .*......*...*...*... | 4 | 0.144140246 | 0.310153846 | chr19:9404893-9404915 | *CTC-325H20.4-ZNF699/CTC-325H20.4* | intergenic |
| TTCTGCCCACGATGACACTGAGG | .*.....*.*......*... | 4 | 0.56142625 | 0.271515151 | chr22:20168160-20168182 | *AC006547.14-XXbac-B444P24.8* | intergenic |
| TACTGACCAGGATGAAGCAGAGG | .....*.*.......*..*. | 4 | 0.049272081 | 0.265306123 | chr2:109253612-109253634 | *LIMS1* | intron |
| AAATGCCTAGGAGGAAGCTGGGG | *.*.........*..*.... | 4 | 0.103917498 | 0.226086956 | chr5:68026266-68026288 | *CTC-537E7.2-CTC-340D7.1* | intergenic |
| AACTTCCGAGGATGAAGCTGGGG | *...*..*.......*.... | 4 | 0.272333333 | 0.22 | chr6:100551945-100551967 | *MCHR2-AS1-PRDX2P4* | intergenic |
| TTCTTCCCAGGATGACGCTAAGG | .*..*..*...........* | 4 | 0.697447183 | 0.204545454 | chr15:27606535-27606557 | *GABRG3-AC144833.1* | intergenic |
| AACTGCCTAAAATGTCGCTGAGG | *........**...*..... | 4 | 0.205917821 | 0.186666667 | chr10:101526549-101526571 | *CUTC-ABCC2* | intergenic |
| TACAGCCCTGGATGGCGCTGGGG | ...*...**.....*..... | 4 | 0.234278012 | 0.185714286 | chr5:178012701-178012723 | *COL23A1* | intron |

| | | | | | | | |
|---|---|---|---|---|---|---|---|
| AACTCCTTAGGATGACGCAGAGG | *...*.*...........*. | 4 | 0.359837588 | 0.182397959 | chr12:26013113-26013135 | *RP11-443N24.3-RP11-443N24.4* | intergenic |
| TACCGCCCAGGCTGAAGCTGGGG | ...*...*...*...*.... | 4 | 0.127203797 | 0.170278638 | chr3:46753357-46753379 | *TMIE-PRSS50* | intergenic |
| TACTGCCGATGAGGAAGCTGAGG | .......*.*..*..*.... | 4 | 0.083678408 | 0.155434782 | chr3:612765-612787 | *AC090044.1* | intron |
| TACTGCCGATGAGGAAGCTGAGG | .......*.*..*..*.... | 4 | 0.083678408 | 0.155434782 | chr21:9539436-9539458 | *Gap-CR381670.1* | intergenic |
| AATTGCTTAGGATGACTCTGGGG | *.*...*.........*... | 4 | 0.410516581 | 0.139648438 | chr3:72514258-72514280 | *RYBP-RP11-654C22.2* | intergenic |
| TTCTGCCGAGAAGGACGCTGAGG | .*.....*..*.*....... | 4 | 0.307297816 | 0.139130435 | chr7:152536523-152536545 | *ACTR3B* | intron |
| TTCTGCCGAGAAGGACGCTGAGG | .*.....*..*.*....... | 4 | 0.307297816 | 0.139130435 | chr7:149967512-149967534 | *ACTR3C* | intron |
| CCCTGCCTGGGATGATGCTGTGG | **......*......*.... | 4 | 0.166395667 | 0.138147567 | chr4:154720444-154720466 | *SFRP2-AC020703.1* | intergenic |
| CACTGCCTAGGCTGGCACTGAGG | *.........*..*.*.... | 4 | 0.08037722 | 0.136842105 | chr13:25476200-25476222 | *CENPJ* | intron |
| TCCTGCCCAGGAAGACCCTGAGG | .*.....*....*...*... | 4 | 0.23590875 | 0.127989658 | chr19:31454160-31454182 | *CTC-400I9.3-AC020952.1* | intergenic |
| TACTGCCTAGGGTTGAGCTGGGG | ...........*.***.... | 4 | 0.004409678 | 0.125185185 | chr5:172376398-172376420 | *ERGIC1* | intron |
| TGCTGCCTAAGAAGACCCTGAGG | .*.......*..*...*... | 4 | 0.217271959 | 0.121628959 | chr11:10208741-10208763 | *RP11-748C4.1-SBF2* | intergenic |
| TACTGCCCTGGATGGTGCTGGGG | .......**.....**.... | 4 | 0.038443137 | 0.12 | chr15:31091096-31091118 | *GOLGA8UP* | intron |
| TACTGCCTTAGAAGATGCTGAGG | ........**..*..*.... | 4 | 0.051127507 | 0.119289941 | chr7:69367849-69367871 | *AUTS2* | intron |
| CCCTGCCTGGGAGGACGCTGCGG | **......*...*....... | 4 | 0.355433782 | 0.117125111 | chr1:2066579-2066601 | *PRKCZ* | intron |
| TAGTGCCTAAGTTAACGCTGCGG | ..*......*.*.*...... | 4 | 0.095245275 | 0.116666667 | chr4:155161554-155161576 | *DCHS2* | intron |
| CACAGCCTAGGATGCAGCTGGGG | *..*..........**.... | 4 | 0.072985333 | 0.111317254 | chr18:72955041-72955063 | *TSHZ1* | intron |
| TACTGGCTAGGATGAACCTAGGG | .....*.........**..* | 4 | 0.025112337 | 0.110294118 | chr6:119236695-119236717 | *MCM9* | intron |
| TACTGCCTCAGATGGTGCTGGGG | ........**....**.... | 4 | 0.035406129 | 0.106666667 | chr19:1908581-1908603 | *ADAT3/SCAMP4* | intron |
| CGCTGCCCAGGATGAGGCTGGGG | **.....*.......*.... | 4 | 0.272333333 | 0.105494506 | chr20:57195347-57195369 | *APCDD1L-AS1-MGC4294* | intergenic |
| CACTGCCTGGGCAGACGCTGTGG | *.......*..**....... | 4 | 0.174873421 | 0.10410642 | chr3:129307423-129307445 | *PLXND1* | intron |
| TCCTGCCAAGGATGGCCCTGGGG | .*.....*......*.*... | 4 | 0.163368333 | 0.096134454 | chr10:133008610-133008632 | *TCERG1L* | intron |
| TACTGCCAGGGATGACCCTTGGG | .......**.......*..* | 4 | 0.147449913 | 0.087843137 | chr5:43246184-43246206 | *NIM1K* | intron |
| TATTACTTAGGATGAGGCTGGGG | ..*.*.*........*.... | 4 | 0.174113559 | 0.074479167 | chrX:71562857-71562879 | *HDAC8* | intron |
| TAAAGCCTAGGAAGACTCTGAGG | ..**........*...*... | 4 | 0.220828507 | 0.071428571 | chr6:22433745-22433767 | *RP3-404K8.2-HDGFL1* | intergenic |
| TACTGCTTGTGATGAGGCTGGGG | ......*.**.....*.... | 4 | 0.094581352 | 0.067708333 | chr6:150559142-150559164 | *PPP1R14C* | intron |
| TAATGACTAGGATGAGGATGTGG | ..*..*.........*.*.. | 4 | 0.031841181 | 0.066666667 | chr16:74008324-74008346 | *RPSAP56-AC009120.4* | intergenic |
| TACCTCCTAGGACCACGCTGGGG | ...**.......**...... | 4 | 0.082499774 | 0.065678903 | chr2:175112699-175112721 | *OLA1* | intron |
| CACTGTCTAGGATTATGCTGAGG | *....*.......*.*.... | 4 | 0.024549488 | 0.065306123 | chr1:65594559-65594581 | *MRPS21P1-AK4* | intergenic |
| TCCTGCCTAGAATGAGGATGTGG | .*........*....*.*.. | 4 | 0.02962442 | 0.065088757 | chr2:163125807-163125829 | *IFIH1* | intron |
| TACAGCCTAAGATCATGCTGGGG | ...*.....*...*.*.... | 4 | 0.035479776 | 0.058608059 | chr1:186968564-186968586 | *PLA2G4A-LINC01036* | intergenic |
| TACTGCATAGGATTATACTGGGG | ......*......*.**... | 4 | 0.009641657 | 0.057435898 | chr6:44481606-44481628 | *CDC5L-RP3-449H6.1* | intergenic |
| TTCTGCCTAGGCTCGCGCTGCGG | .*.........*.**..... | 4 | 0.029531989 | 0.053315106 | chr1:2144990-2145012 | *AL590822.1/RP11-181G12.5/C1orf86/RP11-181G12.4* | intergenic |
| TACTGCTAAGCATGAGGCTGGGG | ......**..*....*.... | 4 | 0.093281959 | 0.042857143 | chr9:97724698-97724720 | *C9orf3* | intron |
| TTCTTCCTAGGATAACTCTGAGG | .*..*........*..*... | 4 | 0.090827917 | 0.040909091 | chr3:18178080-18178102 | *AC132807.1-SATB1* | intergenic |
| TACTTCCTAGGACGATGATGAGG | ....*.......*..*.*.. | 4 | 0.019611103 | 0.039240112 | chr11:17629090-17629112 | *OTOG* | intron |
| TACTGCCTCTGATGCTGCTGAGG | ........**....**.... | 4 | 0.035406129 | 0.038961039 | chr11:58632351-58632373 | *GLYATL2-GLYATL1P2* | intergenic |
| TGCTGCCTGGGAGGACTCTGGGG | .*......*...*...*... | 4 | 0.144140246 | 0.034782609 | chr11:2174136-2174158 | *INS-IGF2/IGF2-INS-IGF2* | intergenic |
| TACTGCCTCTGCTGCCGCTGTGG | ........**.*..*..... | 4 | 0.101277997 | 0.033321941 | chr17:63533165-63533187 | *AXIN2* | **exon** |
| TTCTGCCTAGGCTGGCGCGGTGG | .*.........*..*...*. | 4 | 0.06576318 | 0.031100478 | chr12:43572807-43572829 | *RP11-118A3.1-AC079603.1* | intergenic |
| TTCTGCCCAGGCTGAGGCTGAGG | .*.....*...*...*.... | 4 | 0.127203797 | 0.02944424 | chr1:200862759-200862781 | *C1orf106* | intron |
| TACTGACGAGGCTGAGGCTGAGG | .....*.*...*...*.... | 4 | 0.073249464 | 0.027568922 | chr4:170845432-170845454 | *RP11-205M3.3* | intron |
| AGCTGCCTAGGATGACCGTGTGG | **..............**.. | 4 | 0.119478333 | 0.025098039 | chr17:55538633-55538655 | *MSI2-RP11-118E18.2* | intergenic |
| AACTGCCTGGGATGACTGTGTGG | *.......*.......**.. | 4 | 0.073001262 | 0.022222222 | chr2:220099746-220099768 | *ANKZF1* | **exon** |
| TACTGCCTGGGGTGAGTCTGTGG | ........*..*...**... | 4 | 0.027171265 | 0.018518519 | chr11:79067067-79067089 | *TENM4-MIR708* | intergenic |
| TACTTCCAAGGATTACCCTGAGG | ....*..*.....*..*... | 4 | 0.086229035 | 0.015058824 | chr9:79421778-79421800 | *PRUNE2* | intron |
| TACTCCCTAGGAGGACCCAGAGG | ....*.......*...*.*. | 4 | 0.070548661 | 0.013779425 | chr3:94216126-94216148 | *NSUN3-ARMC10P1* | intergenic |
| GACTGCCTGGGATGAGGGTGCGG | *.......*......*.*.. | 4 | 0.032613551 | 0.013080639 | chr3:193446631-193446653 | *OPA1-RN7SL447P* | intergenic |
| TACTGTCTAGGATTAGCCTGGGG | .....*.......*.**... | 4 | 0.00854056 | 0.008963585 | chr1:217346142-217346164 | *ESRRG-GPATCH2* | intergenic |
| TTCTGCCTAGGAGGAGGCAGAGG | .*..........*..*..*. | 4 | 0.033198795 | 0.008339487 | chr12:40117882-40117904 | *C12orf40* | intron |
